# The lesser Pacific striped octopus, *Octopus chierchiae*: an emerging laboratory model for the study of octopuses

**DOI:** 10.1101/2021.08.04.455148

**Authors:** Anik G. Grearson, Alison Dugan, Taylor Sakmar, Gül Dölen, David H. Gire, Dominic M. Sivitilli, Cristopher M. Niell, Roy L. Caldwell, Z. Yan Wang, Bret Grasse

## Abstract

Cephalopods have the potential to become useful experimental models in various fields of science, particularly in neuroscience, physiology, and behavior. Their complex nervous systems, intricate color- and texture-changing body patterns, and problem-solving abilities have attracted the attention of the biological research community, while the high growth rates and short life cycles of some species render them suitable for laboratory culture. *Octopus chierchiae* is a small octopus native to the central Pacific coast of North America whose predictable reproduction, short time to maturity, small adult size, and ability to lay multiple egg clutches (iteroparity) make this species ideally suited to laboratory culture. Here we describe novel methods for culture of *O. chierchiae,* with emphasis on enclosure designs, feeding regimes, and breeding management. Our results demonstrate the feasibility of multigenerational culture of *O. chierchiae.* Specifically, *O. chierchiae* bred in the laboratory grows from a 3.5-millimeter mantle length at hatching to an adult mantle length of approximately 20-30 millimeters in 250-300 days, with 14-15% survivorship to over 400 days of age in first and second generations. *O. chierchiae* sexually matures at around an estimated six months of age and, unlike most octopus species, can lay multiple clutches of eggs, approximately every 30-90 days. Eggs are large and hatchlings emerge as direct developing octopuses. Based on these results, we propose that *O. chierchiae* possesses both the practical and biological features needed for a model octopus that can be cultured repeatedly to address a wide range of fundamental biological questions.

## 1. Introduction

Cephalopods (octopuses, squid, cuttlefish, and nautiluses) are a diverse and wide-spread class of marine molluscs that have long been useful for scientific research. Studies of their complex nervous systems (Young, 1971), dynamic skin patterns (Hanlon et al., 2011), and arm biomechanics (Gutfreund et al., 2006; Sumbre et al., 2005; Tramacere et al., 2014), have yielded many foundational insights into physiology, developmental biology, and neurobiology (Shigeno and Ragsdale, 2015; Stubbs and Stubbs, 2016; Di Cosmo et al., 2018; Edsinger and Dölen, 2018; Wang and Ragsdale, 2018; Geffeney et al., 2019; Juárez et al., 2019; Maiole et al., 2019; van Giesen et al., 2020; Schnell et al., 2021). Unique amongst coleoid cephalopods, octopuses demonstrate higher-order cognitive capabilities most commonly attributed to vertebrates, including tool use, problem solving, observational learning, and episodic memory (Hochner et al., 2006; Huffard, 2013; Hanlon and Messenger, 2018).

Despite the immense potential for comparative biological research between cephalopods and vertebrates, several biological and technical reasons have hindered the advancement of octopus culture in laboratory settings:

### 1) Water quality

Cephalopods produce a large amount of nitrogenous waste compared to other marine organisms (Boucher-Rodoni and Mangold, 1995). Octopuses have a rapid metabolism and a high rate of waste production (Hanlon, 1987; O’Dor, 1987; Lee et al., 1995). Changes in water parameters, such as high ammonia or nitrogen levels, can greatly impact octopus development and health (Oestmann et al., 1997; Lee et al., 1998). Maintaining optimal water quality at all times is paramount to the success of octopus culture (Boletzky and Hanlon, 1983; Hanlon and Forsythe, 1985; Vidal and Boletzky, 2014).

### 2) Diet

Octopuses are active predators and have specific dietary requirements to maintain rapid growth, especially as hatchlings (Lee et al., 1998; Navarro et al., 2014; Vidal et al., 2014); they require a high protein and relatively low high-quality lipid diet (Miliou et al., 2005; Rosas et al., 2013; Vidal et al., 2014).

### 3) Safe containment

Most octopus species are highly cannibalistic (Ibánez and Keyl, 2010) and solitary as adults; conspecific aggression is very common. Therefore, individual enclosures are required to house animals safely. This increases the space, materials, and amount of seawater necessary for rearing octopuses in the lab. Further, octopuses are notoriously skilled at escaping containment, so the design of octopus enclosures must include sufficient measures to prevent their escape (Wood and Anderson, 2004).

### 4) Life history attributes

Most cephalopod species are semelparous; they die after one reproductive event (Mangold, 1987; Cortez et al., 1995). For example, females of commonly studied octopus species, including *Octopus bimaculoides*, *Octopus vuglaris*, and *Octopus maya*, lay only one clutch of eggs in their lives. Octopus hatchlings can undergo one of two mechanisms of development: indirect and direct development. Small egg species, such as *O. vulgaris*, undergo indirect development; they lay large numbers of small eggs that develop into planktonic paralarvae before undergoing metamorphosis to achieve the benthic stage (Vidal et al., 2014), conditions which are extremely difficult to replicate in the lab. Large egg species, such as *O. bimaculoides*, undergo direct development and hatchlings emerge as fully developed benthic octopuses, which also experience high rates of mortality (Itami et al., 1963; Iglesias et al., 2000; Iglesias et al., 2004; Vaz-Pires et al., 2004; Vidal et al., 2014; Garrido et al., 2018). These natural life history attributes greatly limit octopus reproduction and the rearing of hatchlings in laboratory settings, and multigenerational culture of these species has not yet been achieved (Iglesias et al., 2007).

Here, we overcome these challenges by advancing the laboratory aquaculture of the lesser Pacific striped octopus, *Octopus chierchiae* (Figure 1). *O. chierchiae* is native to the central Pacific coast of the Americas (Voss, 1971) and lives in low intertidal zones at water depths up to 40 meters (Rodaniche, 1984; Hofmeister et al., 2011). Several unique biological features make *O. chierchiae* particularly amenable for laboratory culture and scientific research. As adults, *O. chierchiae* are very small: the largest *O. chierchiae* adult mantle length previously reported was 25 millimeters (mm) (Rodaniche, 1984). Other octopus study species, such as *O. bimaculoides, O. maya,* and *O. vulgaris*, can reach a mantle length three to ten times larger than *O. chierchiae* (Jereb et al., 2014). Unlike most octopus species, *O. chierchiae* is iteroparous. Individuals can undergo multiple rounds of reproduction and lay successive clutches of eggs before death, which is useful for planning and managing the reproductive outputs of a colony of research animals (Rodaniche, 1984; Hofmeister et al., 2011). *O. chierchiae* is a large egg species and lacks a paralarval phase (Rodaniche, 1984); hatchlings are direct developing and emerge with a highly developed nervous system and innate hunting behaviors (Vidal et al., 2014). Mature males and females are visually and behaviorally sexually-dimorphic: the tip of the third right arm in male *O. chierchiae* lacks suckers and is modified with a hectocotylus, a smooth, hooked organ used to pass spermatophores to the female during mating (Voss, 1968; Rodaniche, 1984; Figure 1B). Males also exhibit a visually striking arm-twirling behavior in which they rapidly shake the tips of their arms (Hofmeister et al., 2011), which here we call “tasseling”. In addition, *O. chierchiae* are marked by sharply contrasting black and white striping patterns (Figure 1A), which are unique to each individual and can allow for non-invasive identification (Hofmeister et al., 2011). Thus, culture of *O. chierchiae* requires significantly less physical space to produce many generations of directly developing offspring that can be visually identified, making them particularly advantageous for laboratory study as compared to other octopus species used in research.

**Figure 1.**
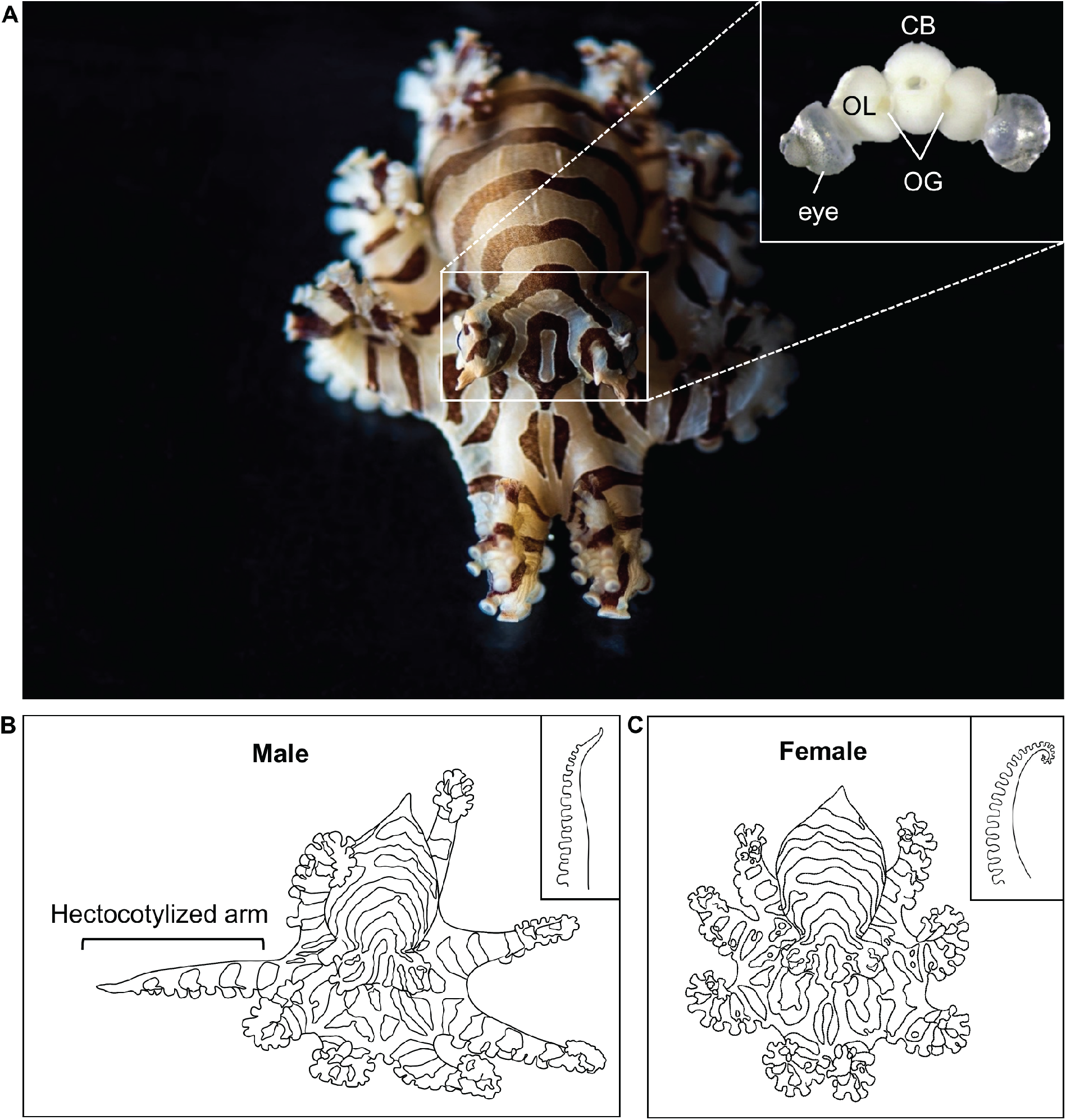
Adult Octopus chierchiae. (A) Dorsolateral view of an adult *O. chierchiae,* an iteroparous species of octopus native to the Pacific coast of North and Central America. Visual stripes are present 7-14 days after hatching and can be used to distinguish individuals. Inset depicts central nervous system with eyes attached. Two large, kidney-shaped optic lobes (OL) flank the central brain mass (CB; supraesophageal and subesophageal masses). The optic glands (OG) sit on top of the optic tract between the OL and the CB. (B) Illustration of a male *O. chierchiae* with the hectotoyllzed arm extended. Male adult *O. chierchiae* possess a specialized organ on the tip of the third right arm, the hectocotylus (depicted in inset), which has a smooth, sucker-less, hook-like appearance and is used to pass spermatophores to the female during mating. (C) Illustration of a female O. *chierchiae,* lacking a hectocotylized arm. Female adult *O. chierchiae* possess suckers along the full length of all arms (depicted in inset). Image A courtesy of Tim Briggs.

Culture of *O. chierchiae* has been attempted multiple times since the 1970’s without success. The primary reasons for failure include inadequate diet and nutrition at early life stages, inadequate holding enclosures and seawater systems, and gaps in general knowledge of their biology and behavior (Rodaniche, 1984). Collection of this species has been infrequent and challenging since the 1970s, so little is known about *O. chierchiae* natural history in the wild. This makes the ongoing culture of this species critical for laboratory use and for direct observation of their behaviors. Multi-generational laboratory culture of octopuses has only been reported in one other species, *O. maya* (Van Heukelem, 1977). Here, we document the first successful multi-generational culture of *O. chierchiae* at the Marine Biological Laboratory in Woods Hole, MA. Through the creation of new low-cost enclosures using commonly sourced and commercially available materials, an amended diet, and rigorous cleaning and maintenance protocols, we successfully raised *O. chierchiae* in the laboratory for three generations. Octopuses grew from a 3.5 mm mantle length at hatching to an adult mantle length of approximately 20-30 mm in 250-300 days, with 14-15% survivorship to over 400 days of age in first and second generations. We estimate that *O. chierchiae* sexually mature at around six months of age, and can lay multiple clutches of eggs, approximately every 30-90 days. Based on these results, we propose that *O. chierchiae* possesses both the practical and biological features needed for a model octopus that can be cultured repeatedly to address a wide range of fundamental biological questions.

## 2. Methods

### 2.1 Water chemistry

Water quality was tested weekly, and all water parameters were kept within optimal ranges as outlined in previous reports of cephalopod culture (Hanlon and Forsythe, 1985; Grasse, 2014). pH was tested using an Orion Star A221 meter and maintained between 8.0-8.3 using Two Little Fishes Kalkwasser to raise pH values when needed. Seawater temperature was tested using an Orion Star A221 meter and maintained between 72°F-76°F (22°C-24°C) using 500w Finnex immersion heaters. Salinity was kept between 33-35 parts per thousand (ppt), tested using a Hach Hq40d multi meter. Salinity was lowered by adding reverse osmosis (RO)/deionized (DI) water or raised by adding hypersaline Marine Enterprise Crystal Sea Bioassay Laboratory Formula salt mix. Dissolved oxygen was tested using a Hach Hq40d and maintained at >80% through water changes and Bubblemac Industry air stones. Ammonia and nitrite levels were measured using a Hach DR3900 spectrophotometer and were maintained at 0 parts per million (ppm) and nitrate levels were kept at <15 ppm, via water changes and water purification methods described below.

### 2.2 System design and tray maintenance

Two types of seawater systems were used during the culture of this species: closed and semi-closed (Figure 2). Both system types produced successful results for keeping and culturing *O. chierchiae*. Closed refers to a system with a reservoir of artificial seawater that recirculates continuously through the appropriate filtration (mechanical, chemical, and biological) and ultraviolet (UV) sterilization. The artificial seawater was made by mixing Marine Enterprise Crystal Sea Bioassay Laboratory Formula with RO/DI water, and 10% water changes were performed weekly as needed depending on water quality. Due to fluctuations in natural water chemistry in Vineyard Sound throughout the year, closed systems were beneficial in supplying consistent water chemistry and conditions.

**Figure 2.**
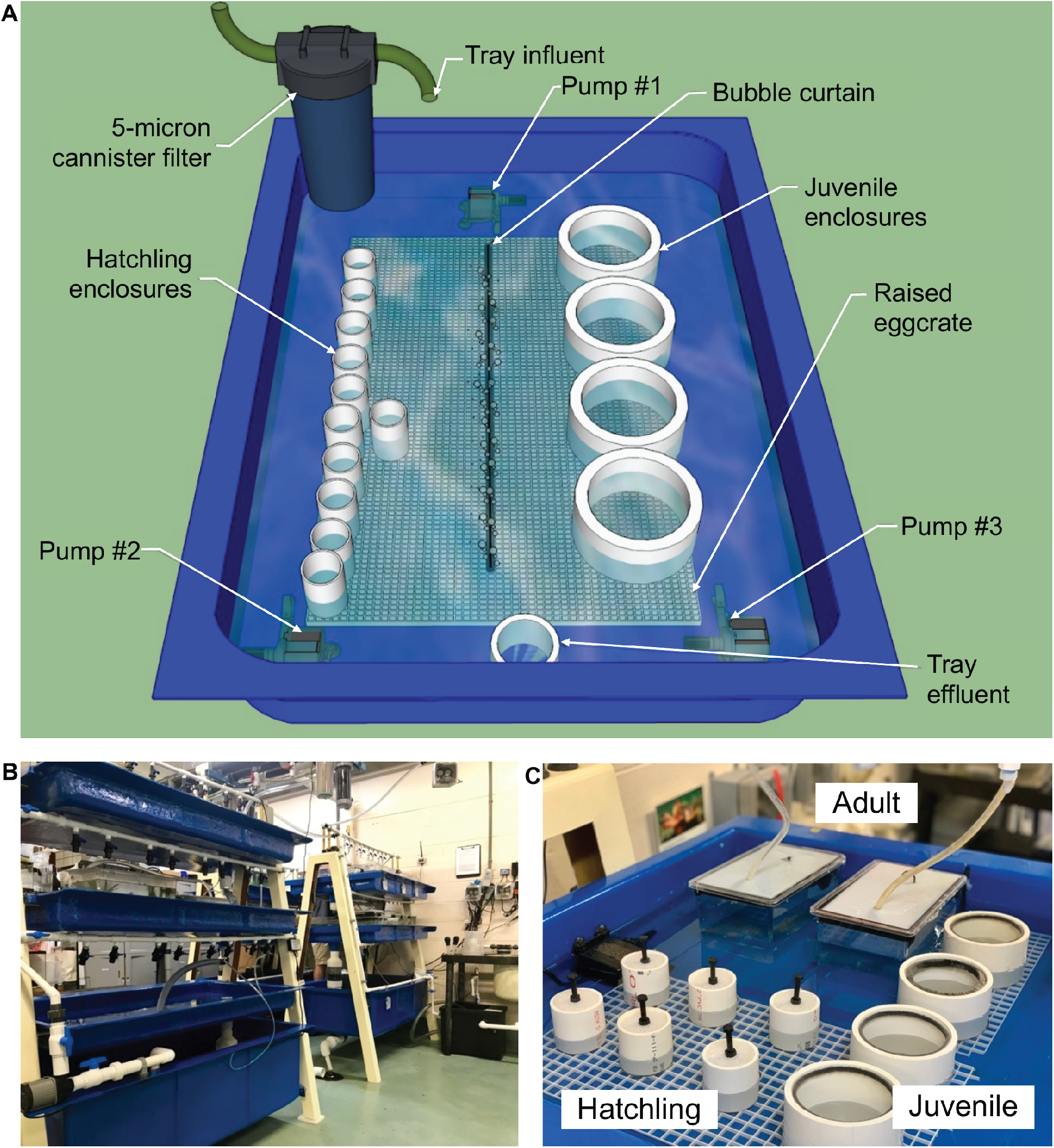
System and enclosure design. (A) Tray design used at the Marine Biological Laboratory. (B) A-frame system design made of fiberglass supports with a large fiberglass reservoir on the bottom and two shallow fiberglass trays above. Water circulates from the reservoir up to the trays via manifolds and discharges back down to the reservoir. Seawater recirculates through the appropriate filtration (mechanical, chemical, and biological) and ultraviolet (UV) sterilization. (C) Adult, juvenile, and hatchling enclosure designs.

Semi-closed seawater systems consisted of filtered seawater from Vineyard Sound, MA, USA with a turnover rate of 200% per day and the seawater recirculating through the appropriate filtration (mechanical, chemical, and biological) and UV sterilization. Prior to entering the systems, the seawater was filtered via a Stark horizontal sand filter consisting of three grades of Dryden Activated Filter Media^®^ and topped with granulated activated carbon. Seawater was then directed through an 80W Emperor Aquatics SMART UV Sterilizer, 50-micron pleated cartridge filter, and granulated activated carbon cartridge and Seachem Cuprisorb™ packet.

Both closed and semi-closed systems used aerated, submerged Kaldnes K1 media for biological filtration. Size and wattage of UV sterilizers were dictated by the water volume (100-150 gallons) and flow rates of each specific system (~90 ml/second in fiberglass trays). Submerged granulated activated carbon was used in each reservoir and replaced monthly. All seawater was mechanically filtered via 50-micron bag filters that were changed weekly. Protein skimmers were used in the reservoirs where appropriate and sized according to each system volume. Protein skimmers were cleaned two to three times weekly, the bag filters once weekly, the cartridge filters every two weeks, and the entire trays once every one to two months. These were scrubbed and/or sprayed clean and rinsed with tap water to remove detritus, soaked in 10% bleach/water solution, then soaked in a mixture of 10% sodium thiosulfate (Proline™)/ water solution to neutralize the bleach, and finally rinsed thoroughly with DI water before re-use. All systems were kept on a 12-hour light/dark cycle.

Both closed and semi-closed systems utilized custom-built A-frame fiberglass supports with a large fiberglass reservoir on the bottom and two shallow fiberglass trays above (Figure 2B). Water circulated from the reservoir up to the trays via manifolds and discharged back down to the reservoir. All individuals were held within the shallow trays of each seawater system, with three to four inches of water depth (Figure 2A). In hatchling and juvenile enclosures (described in Section 2.3), seawater was exchanged through a fine mesh bottom in each enclosure (Figure 2B). Hatchling and juvenile enclosures were elevated off the tray bottom using ½-inch by ½-inch square-cell polystyrene egg crate tables to support seawater exchange through each enclosure’s mesh bottom. Seawater exchange in enclosures was further increased by circulating the water in the tray with small submersible pumps (Rio) in each corner and a bubble curtain running lengthwise down the tray (Figure 2A). Adult enclosures received new seawater directly from valve-controlled supply manifolds on each system (Figure 2B). Displaced effluent water exited through a screened bulkhead back to the fiberglass tray. All seawater in contact with non-native organisms at Marine Biological Laboratory was sterilized with ozonation before returning to the ocean.

### 2.3 Enclosure design and maintenance

Three enclosure designs were used based on the age and size of the octopus (Figure 2C). After hatching, each *O. chierchiae* was housed in a separate enclosure. Once an octopus grew to about half the size of its enclosure (i.e., opposite arm tips reached approximately half of the enclosure diameter), it was transferred to a new enclosure of the next size. The approximate ages at which we moved animals is indicated below. All enclosures were provided with 1-2 dens (see Supplementary Figure S1), and juvenile and adult enclosures also included shells and rocks of varying sizes for enrichment. Dirty enclosures, dens, and materials were scrubbed, cleaned, and rinsed using the cleaning methods described in Section 2.2.

#### Hatchling enclosures

hatchlings were housed in 1¼-inch polyvinyl chloride (PVC) enclosures with a 125-micron polyester mesh bottom and were provided ⅛-inch or ¼-inch barbed fittings as dens. Hatchlings readily escape their enclosures, which can lead to mortality due to desiccation or becoming lost in the system tray. Tight-fitting lids are thus critical to minimize hatchling mortalities due to escape. The lids were removed during feeding, and extra attention was taken to note the location of the octopus during feeds and to ensure the lid was carefully placed back onto the enclosure without crushing the octopus. Hatchlings are very sensitive to changes in water parameters, physical disturbances, or the growth of foreign organisms in their enclosures such as algae, hydroids, or cyanobacteria. To prevent unwanted growth, dirty enclosures, dens, and enrichment materials were replaced with clean ones once every two weeks.

#### Juvenile enclosures

at approximately three months of age, octopuses were moved to 4-inch PVC enclosures with an 840-micron polyester mesh bottom and were provided with 90° PVC elbows as dens. To deter octopuses from escaping out of the juvenile enclosures, we attached adhesive Velcro (rough hook side exposed) or adhesive weatherstrip window pile to the top-inner rim of the PVC tube. These materials are difficult for octopus suckers to adhere to and successfully prevented escape. Juvenile enclosures were replaced with clean ones once every three weeks.

#### Adult enclosures

at approximately six months of age, octopuses were moved to 2.84 L (23.1×15.2×15.5 cm) plastic aquaria, each fitted with an opaque High-Density Polyethylene (HDPE) weighted lid, seawater supply line, bulkhead, and filter screen allowing for water to flow directly into and out of the enclosure. A 1250-micron polyester mesh was placed within the filter screen of enclosures with egg-brooding females to prevent any potential hatchlings from escaping. Adult enclosures were placed directly in the trays as described in Section 2.2 (Figure 2A) and were provided with various sizes of dens. Adult enclosures were replaced with clean ones once every six to eight weeks.

### 2.4 Collection and procurement of wild specimens

Historically, *O. chierchiae* have been spotted at several locations along the Pacific coast of Central America, with a few reported sightings as far North as the Gulf of California. For a full map of these locations, see Supplementary Figure S2. In 2018 and 2019, seven adult *O. chierchiae (*five females, two males) were purchased from a commercial wholesale vendor, Quality Marine (California, USA), to establish cultures. These were wild collected from the Pacific intertidal zones of Nicaragua.

### 2.5 Mating trials and monitoring egg development

To initiate mating, sexually mature female *O. chierchiae* were presented with males. To the best of our knowledge, no previous studies have reported absolute age of sexual maturity in *O. chierchiae*, so indirect measures were used to determine an octopus’ sexual maturity. Hofmeister et al. (2011) reported an “arm-twirling” behavior by male *O. chierchiae* during mating. As such, incidence of the arm-twirling behavior, which we call “tasseling”, was used as a proxy for sexual maturity in males. In some cases, this behavior was also used to identify the individual as male. Females were deemed to be sexually mature when they reached the age at which males from the same clutch began to display the tasseling behavior, or when they laid their first clutch of eggs (in most cases, this occurred after a mating trial, although some of the wild-collected females laid eggs before being mated in the laboratory) (see Rodaniche (1984)).

Mating trials were performed two to four times per week in one of two methods: 1) introducing the male into the female’s enclosure, or 2) introducing the female followed by the male to a new adult enclosure. In both cases, water flow was turned off and all dens and enrichment materials were removed. No noticeable differences in mating behavior were observed with one method over the other and both were used interchangeably. Mating trials were monitored to ensure the male successfully inserted his hectocotylus into the female’s mantle opening. Once the male and the female separated, which could take up to an hour, individuals were returned to their home enclosures. Females and males were paired based on age, size, how recently that individual had been mated, and whether that individual had previously sired/laid a viable clutch of eggs. Individuals were mated repeatedly until they started showing signs of senescence, including disorientation, inappetence, discoloration, and abnormal locomotion (Anderson et al., 2002). A mating trial was deemed unsuccessful if a female did not lay a viable clutch of eggs within a month of the mating trial, at which point the female could be mated once again, often with a different male.

After a mating trial, females’ dens were visually inspected during daily feeding for presence of eggs. Once a female laid a clutch of eggs, development of the eggs was monitored weekly until hatching and regular enclosure maintenance (described in Section 2.3) was not resumed until after eggs had hatched. The total number of eggs laid was not counted to avoid disturbing incubating females. The number of hatchlings produced per clutch of eggs laid per female was counted and the mean calculated for every generation. A one-way analysis of variance (ANOVA) followed by a pairwise Tukey’s multiple comparison test were performed to compare the means using GraphPad Prism version 8.0.0 for Windows (GraphPad Software, San Diego, California USA, www.graphpad.com).

### 2.6 Feeding

Previous attempts at rearing *O. chierchiae* in captivity used live brine shrimp as the main food source, to little success (Rodaniche 1984). We tested a variety of diets for *O. chierchiae*, including live copepods, frozen shrimp, frozen fish, live grass shrimp, and live crabs. In response to all but the last two food sources, *O. chierchiae* exhibited no interest in eating (i.e., rejected the food item) and showed increased mortality.

In our study, live-collected grass shrimp (*Paelomnetes*, ~ 38 mm from head to tail) from marine estuaries near Woods Hole, MA and crabs (*Hemigrapsus sanguineus,* 35-40 mm in carapace width) from rocky beaches near Woods Hole, MA were offered to *O. chierchiae* to consume *ad libitum*. Hatchlings were fed 1/5^th^ of a freshly killed grass shrimp twice daily (morning and afternoon). Individuals in juvenile enclosures were fed ½ of a freshly killed grass shrimp once daily in the afternoon and leftover food was removed the next morning. Adult males and non-egg-brooding females were fed a single live crab with the claws removed (to reduce the risk of the prey harming the octopus) once in the afternoon three to four times weekly. Egg-brooding females were hand-fed one whole grass shrimp (*Paelomnetes*) three to four times weekly. Any leftover food, dead or alive, was removed at the subsequent feeding to minimize waste buildup.

### 2.7 Measuring mantle length

To quantify growth in *O. chierchiae,* we measured mantle length (tip of mantle to midpoint between the eyes) of up to five hatchlings per clutch every seven days from hatching until death or the end of the study (see below). Mantle length was measured by photographing the octopus in a clear Pyrex petri dish filled with filtered seawater resting atop a flat six-inch ruler. To minimize stress, we tried to take the photo within a minute of moving the octopus to the dish. We collected data from a total of 66 individual octopuses from the first and second generations. Because few third-generation hatchlings were available, they were excluded from this analysis to reduce the risk of stress caused by increased handling. Images were imported to ImageJ for calibration and mantle measurement. Pooled averages of mantle length were taken at seven-day increments (i.e., age 0-7 days, 7-14 days, 14-21 days, and so on). In early 2020, the COVID-19 pandemic impeded the collection of growth data, and weekly measurements were terminated. At this point in time, an additional 40 adult octopuses (i.e., > 280 days old) were randomly selected for mantle length measurement.

### 2.8 Recording mortalities and survival analysis

Cause of mortality and date were recorded at time of death. Natural deaths were defined as any mortality not caused by direct human interference or error and included death due to old age or any unknown health-related issues. Other non-natural causes of death included being crushed under enclosure lids (this only occurred with the smallest hatchlings), escape from enclosures, euthanasia with 2-5% ethanol after observing signs of senescence or compromised health, use in experimental diets or enclosures, stress caused by external factors (such as poor water quality), system problems, or other unknown causes (i.e., cause of death was not recorded at time of mortality). Available mortality data was pooled, grouped by generation, and used to plot a survival curve of the population by generation. Mean survival in days was calculated for each generation and a one-way ANOVA followed by a pairwise Tukey’s multiple comparison test were performed to compare the means using GraphPad Prism version 8.0.0 for Windows (GraphPad Software, San Diego, California USA, www.graphpad.com). The survival analysis was performed using mortality data from a total of nine first-generation clutches, 15 second-generation clutches, and one third-generation clutch. Individuals whose mortality data was lost or otherwise unavailable (*n*=28) were not included in these analyses.

## 3. Results

### 3.1 *Life history of* O. chierchiae *in laboratory culture*

We cultured three consecutive generations of *O. chierchiae* under laboratory conditions from 2018-2020. From the original seven wild collected adults, a total of nine first-generation clutches, 15 second-generation clutches, and one third-generation clutches were raised to adulthood. These culture efforts produced a total of 771 octopuses: 350 first-generation, 396 second-generation, and 25 third-generation octopuses (Figure 3A; Figure 4D). The overwhelming majority of lab-cultured individuals exhibited normal physical features, with the notable exception of an octopus with 16 arms from a third-generation clutch, which was found dead in its enclosure at four days of age.

**Figure 3.**
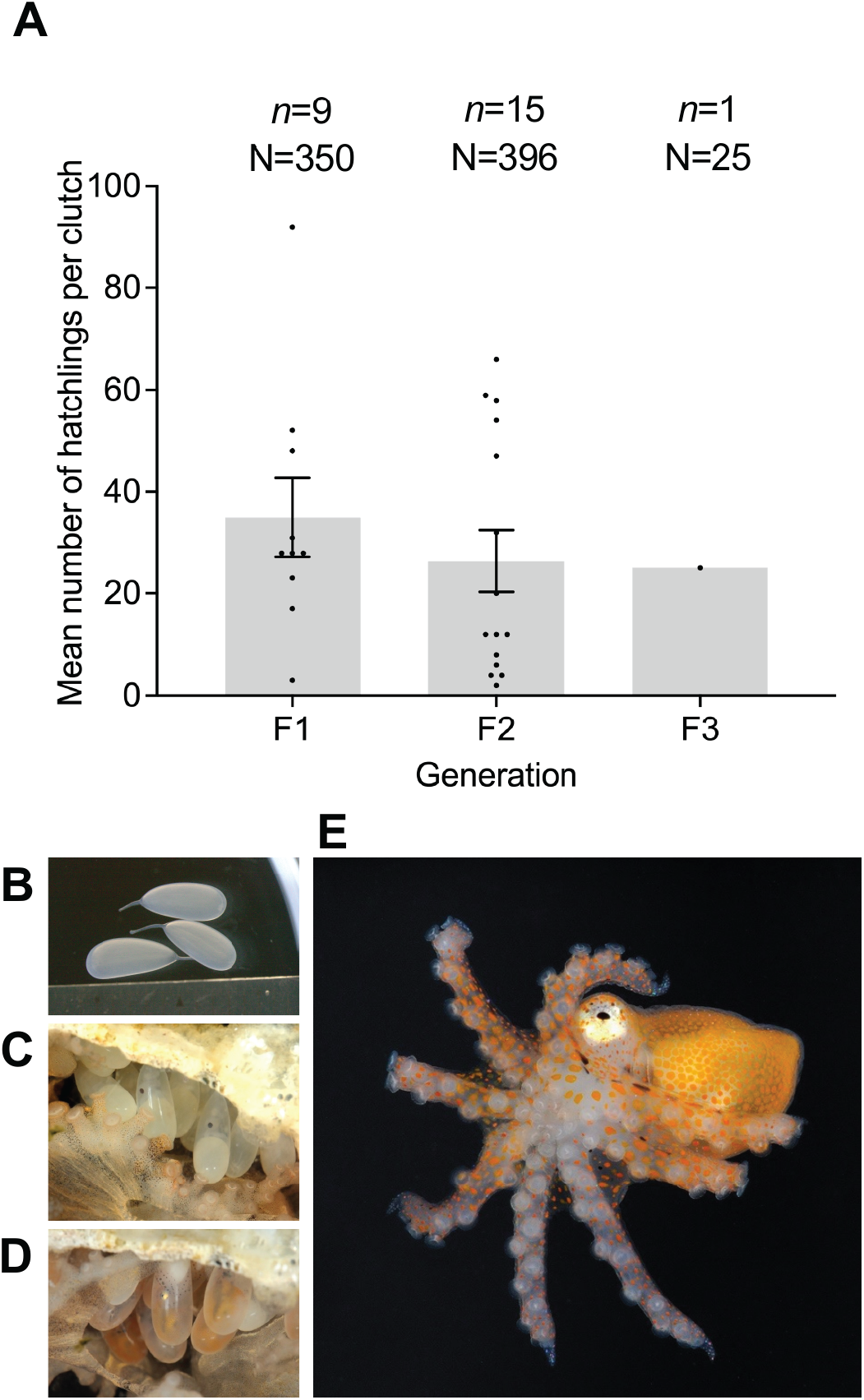
*O. chierchiae* eggs and hatchlings. (A) Mean (±SEM) hatchlings per clutch by generation. A one-way analysis of variance revealed no significant difference between the means of first and second generations (p>0.05). The number of hatchlings for the one third-generation clutch is reported. (B) Translucent, freshly-laid eggs. (C) Eggs with visible eyespots and yolk sac at approximately 17 days post-laying. (D) Eggs with visible chromatophores at approximately 30 days post-laying. (E) O. chierchiae hatchling with bright orange pigmentation; F1=first generation; F2=second generation; F3=third generation; n=number of clutches; N=total number of hatchlings; SEM=standard error of the mean.

**Figure 4.**
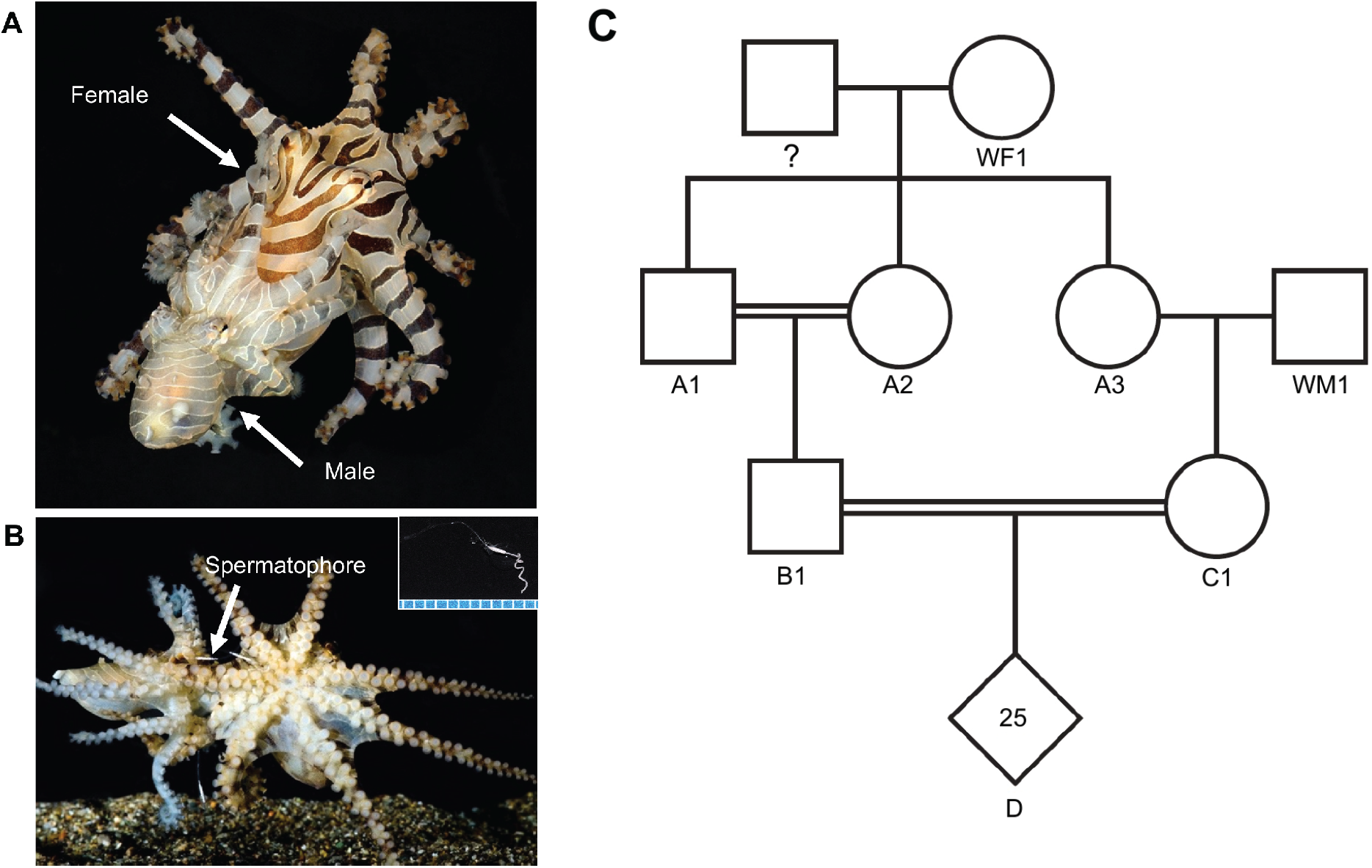
*O. chierchiae* mating and reproduction. **(A)** Male (smaller octopus with pale coloration) and female (larger octopus with dark banded coloration) mating (dorsolateral view). The male inserts the tip of his third right arm, called the hectocotylus, into thefemale’s mantle opening to deliver spermatophores. **(B)** Visible spermatophores during mating. Insetdepictsaspermatophore (scale: millimeters). **(C)** Pedigree showing parental lineage of a third-generation clutch. Squares represent males, circles represent females, diamonds represent clutches, double solid lines highlight inbred pairs. Female WF1 was a wild-collected female. She had mated previously with unknown partner(s)and produced Clutch A upon arrival in the lab. Two offspring from Clutch A (A1 and A2) were bred together and another offspring from Clutch A (A3) was outbred with a wild-collected male (WM1) to produce two second generation clutches. Two individuals from these second-generation clutches (B1 and C1) mated to produce a third-generation clutch (D)with 25 hatchlings. Note Tor clarity, non-reproductive offspring of thefirst-and second-generation clutches are not represented in this pedigree.

Octopuses were moved to juvenile and adult enclosures when they outgrew the size constraints of their given enclosure (i.e., the distance between opposite arm tips reached approximately half of the enclosure diameter). 162 individuals were moved to 4-inch PVC juvenile enclosures at a mean age of 94 days (SEM: 1.8) and 83 individuals were moved to 2.84 L plastic aquaria adult enclosures at a mean age of 211 days (SEM: 3.3) (Table 1).

**Table 1.**
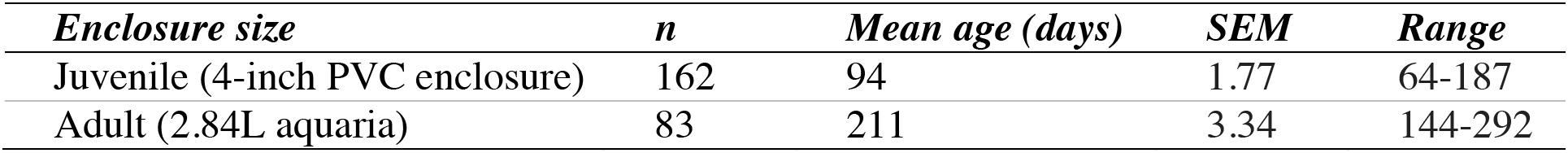
Mean age of *O. chierchiae* when moved to larger enclosures. Octopuses were held in 1¼-inch PVC enclosures from birth until moved to juvenile enclosures. Juvenile enclosures were 4-inch PVC enclosures (see Section 2.3). Adult enclosures were 2.84 L plastic aquaria (see Section 2.3). Octopuses were moved to juvenile and adult enclosures when they outgrew the size constraints of their given enclosure (i.e., the distance between opposite arm tips reached approximately half of the enclosure diameter). *n*=number of individuals; SEM=standard error of the mean.

Both males and females exhibited typical signs of octopus senescence (Anderson et al., 2002) after reaching ages of over a year and a half. These behaviors sometimes included disorientation, inappetence, discoloration, and abnormal locomotion (Supplementary Figure S3). No individuals reproduced after onset of senescence, although senescence did not appear to be triggered by reproduction, as it is in other octopus species (Anderson et al., 2002; Di Cristo, 2013; Wang and Ragsdale, 2018).

### 3.2 Mating and reproduction

A total of ten males were recorded exhibiting the tasseling behavior (see Supplementary Video 1) in their home enclosures (Table 2), which was used as an indication that the male was sexually mature, at a mean age of 176 days (SEM: 8.60). The number of males tasseling was likely much higher as it only began to be recorded in the last year of the study.

**Table 2.** Mean age of *O. chierchiae* at estimated sexual maturity. The tasseling behaviour was used to estimate sexual maturity in males. The first incidence of laying a clutch of eggs was used to estimate sexual maturity in females. *n*=number of individuals; SEM=standard error of the mean.

| *Sexual maturity* | *n* | *Mean age (days)* | *SEM* | *Range* |
| --- | --- | --- | --- | --- |
| Males (tasseling) | 10 | 176 | 8.60 | 136-208 |
| Females (first clutch laid) | 31 | 241 | 4.59 | 196-299 |

We conducted a total of 88 mating trials and 22 octopuses (15 females and 7 males) reproduced. Typically, the female initiated mating by slowly approached the male, who was often exhibiting the tasseling display. This was followed by the male pouncing on the female, grasping her by the mantle from the posterior end, and displaying a pale coloration while the female maintained her dark, banded coloration (Figure 4A; for videos of mating trials, see Supplementary Videos 2 and 3). Beak-to-beak mating, a behavior previously reported in LPSO (Caldwell et al., 2015) and in *O. chierchiae* (Hofmeister et al., 2011), was observed in rare instances and did not lead to egg-laying. Successful mating occurred when the male inserted his hectocotylus into the female’s mantle and delivered spermatophores (Figure 4B-4C), which was visually confirmed in 38/88 mating trials. Occasionally, a female would eject a spermatophore during or after mating. The duration of time between placing both octopuses in the enclosure to their separation after mating was recorded for 43 mating trials, with a mean duration of 56.6 minutes (SEM: 3.79) but ranged from 15 minutes to over an hour.

Females laid eggs in their dens between 14-30 days after successful mating. Before laying a clutch of eggs, a female’s mantle would swell and, in some cases, eggs could be seen through the partially translucent mantle epidermis by visual inspection. 31 females laid their first clutch of eggs at a mean age of 241 days (SEM: 4.59), which was used to estimate approximate age at sexual maturity in females (Table 2). Some females laid eggs before exposure to a male; these eggs were unfertilized and non-viable. A few females were discovered with mature yet unfertilized eggs in their mantle after death at as early as 134 days of age. Eight females raised in the lab laid multiple clutches; four of them were wild-collected and four were from the first generation. In most of these cases, each clutch was the result of a separate mating trial event. Five of these females laid two clutches of eggs and the other three females laid three clutches of eggs. The duration of time between laying subsequent clutches of eggs varied between 30 to 90 days. In one instance, a first-generation female laid two consecutive clutches of eggs after a single mating event, separated by 82 days. Several of the wild females laid a clutch of eggs before having been mated in the laboratory. One of these females laid two clutches of eggs without ever being introduced to a male in the laboratory, separated by 66 days. For full data on reproductive mating pairs, see Supplementary Table S1. These observations align with similar ones reported by Rodaniche (1984), where a single female *O. chierchiae* mated twice and laid three viable clutches of eggs, the last of which was laid 83 days after mating.

While brooding, females remained in their dens and positioned themselves below or next to the eggs. Females continued to feed on the shrimp provided to them while brooding, a behavior that has been previously reported in *O. chierchiae* (Rodaniche, 1984), and evocative of the first stages of maternal care in *O. bimaculoides* (Wang and Ragsdale 2018). Eggs were approximately 6.20 mm in length and were white/translucent in coloration (Figure 3B). Egg development progressed in a similar manner to that of LPSO (Caldwell et al., 2015). At approximately 17 days post-laying, a yolk sac and eyespots could be seen developing in the embryo as two dark spots near the center of the egg (Figure 3C). At approximately 30 days post-laying, bright orange chromatophores could be seen in the developing embryos (Figure 3D). Hatchling octopuses, typically bright orange in coloration (Figure 3E), began emerging from eggs between 40-50 days after laying. The mean number of hatchlings that emerged per clutch was 35.0 (SEM: 7.7) for first-generation clutches (*n*=9), 26.4 (SEM: 6.1) for second-generation clutches (*n*=15), and the total hatchlings for the third-generation clutch was 25 (Figure 3A). A one-way ANOVA and Tukey’s multiple comparisons test revealed that there was no significant difference between the means for first and second generations (*p*>0.05).

Occasionally, females laid non-viable eggs or discarded eggs from their den while brooding, usually fewer than ten eggs. Eggs discarded before embryonic chromatophores were visible did not hatch, while most eggs discarded after formation of embryonic chromatophores hatched if they were stored in a 1½ inch PVC enclosure after being discarded. In four instances, brooding females discarded between 10-20 eggs, and the resulting clutch yielded fewer than five hatchlings. Of these, two of the clutches were the second laid by a female while the other two were laid by one-year-old females from the same first-generation clutch.

Five sexually mature males were mated several times without any observed negative effects to their health. One male in the first generation that survived to 402 days of age was mated eight times (with five different females) and sired eight clutches of eggs, producing a total of 214 viable hatchlings (Supplementary Table S1). Three of these females were siblings from the same clutch of eggs as the male.

### 3.3 Growth

Hatchlings were bright orange (Figure 3E) and stripes began to be visible after approximately 7-14 days. Hatchlings were benthic and direct developing with a mean mantle length of 3.5 mm (SEM: 0.07; *n*=27) during the first week of life (Figure 5). Rodaniche (1984) reported a similar mantle length in laboratory-raised *O. chierchiae* hatchlings, but due to lack of appropriate food, growth remained static after two weeks, and mortality increased. On a diet of *Palaemonetes* shrimp and *H. sanguineus* crabs, preliminary observations in this study suggest that mantle length growth is linear from birth until approximately 150-200 days of age, after which it becomes asymptotic (Figure 5). Variability in average mantle length increased dramatically after ~180 days post-hatch, as many individuals in the growth group died before reaching the end of the study and sample size decreased (Figure 5, Supplementary Figure S4). Based on the data from 40 randomly sampled adults between 280 and 800 days of age, average mantle length did not increase with age after 280 days post hatch (Figure 5). The maximum mantle length recorded was 40 mm in two individuals: one at 550 days post hatch and the other at 799 days post hatch, both of which were from the group of randomly sampled adults.

### 3.4 Survival

Across all three generations, mortality was highest during the first 30-60 days post-hatch. The largest drop in survival occurred in the first five days post-hatch in the third generation (Figure 6A). The mean survival for first, second, and third generations was 128.7 days (SEM: 9.9), 133.3 days (SEM: 8.5), and 25.4 (SEM: 13.7) respectively (Figure 6B). Pair-wise multiple comparisons revealed that mean survival in the third generation was significantly lower than in first and second generations (*p*≤0.05.) and there was no significant difference between survival in first and second generations (Figure 6B).

**Figure 6.**
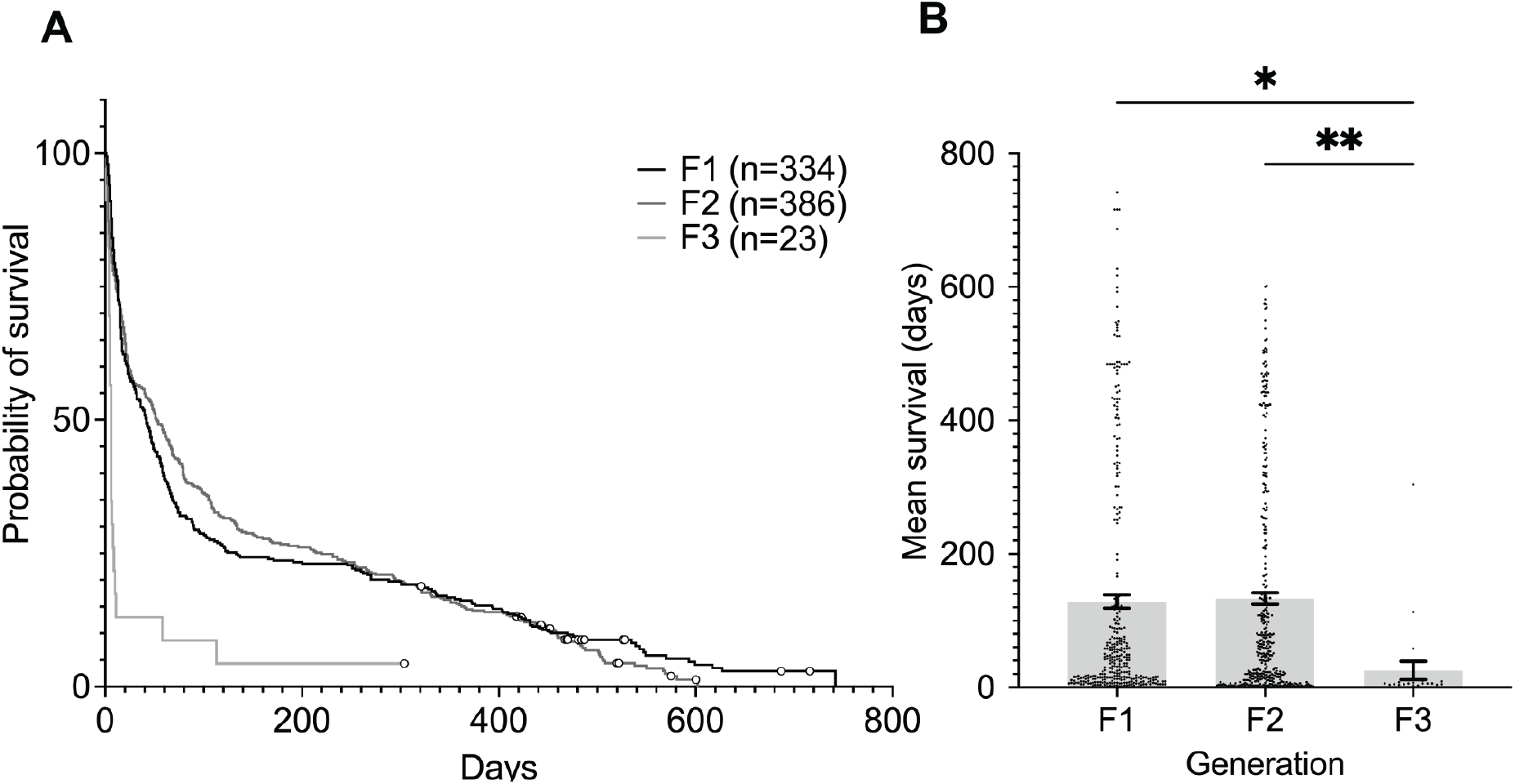
*O. chierchiae* survival by generation. Mortality data was pooled and grouped by generation. Individuals surviving to the end of the study were treated as censored observations, represented by circles. (A) Survival curve by generation. (B) Mean survival (±SEM) in days by generation. A one-way analysis of variance followed by Tukey’s multiple comparison test revealed that mean survival in the first and second generations was significantly higher than in the third generation. F1=first generation; F2=second generation; F3=third generation; n=number of individuals; SEM=standard error of the mean; *=statistically significant at p< 0.05; **=statistically significant at p<0.005.

A total of 50 (15.0%) and 54 (14.0%) individuals survived to 400 days or older in the first- and second-generation clutches, respectively. Of these, 11 (3.3%) survived to >500 days of age and three (0.9%) to >600 days of age in the first generation, while 13 (3.4%) survived to >500 days of age and one (0.3%) to >600 days of age in the second generation. A single individual from the third-generation clutch (4.3%) survived to 387 days of age. For mean proportions of survivors see Supplementary Figure S5. The oldest *O. chierchiae* we recorded in this study was a first-generation individual that survived to 742 days of age.

A total of 315, 367, and 22 mortalities were recorded for first, second, and third generations, respectively. Across all three generations, the largest proportion of mortalities were natural deaths: 54.9%, 76.6%, and 100% for first, second, and third generations, respectively (Table 3). The proportion of mortalities due to human error was much higher in the first generation compared to subsequent generations (Table 3).

**Table 3.**
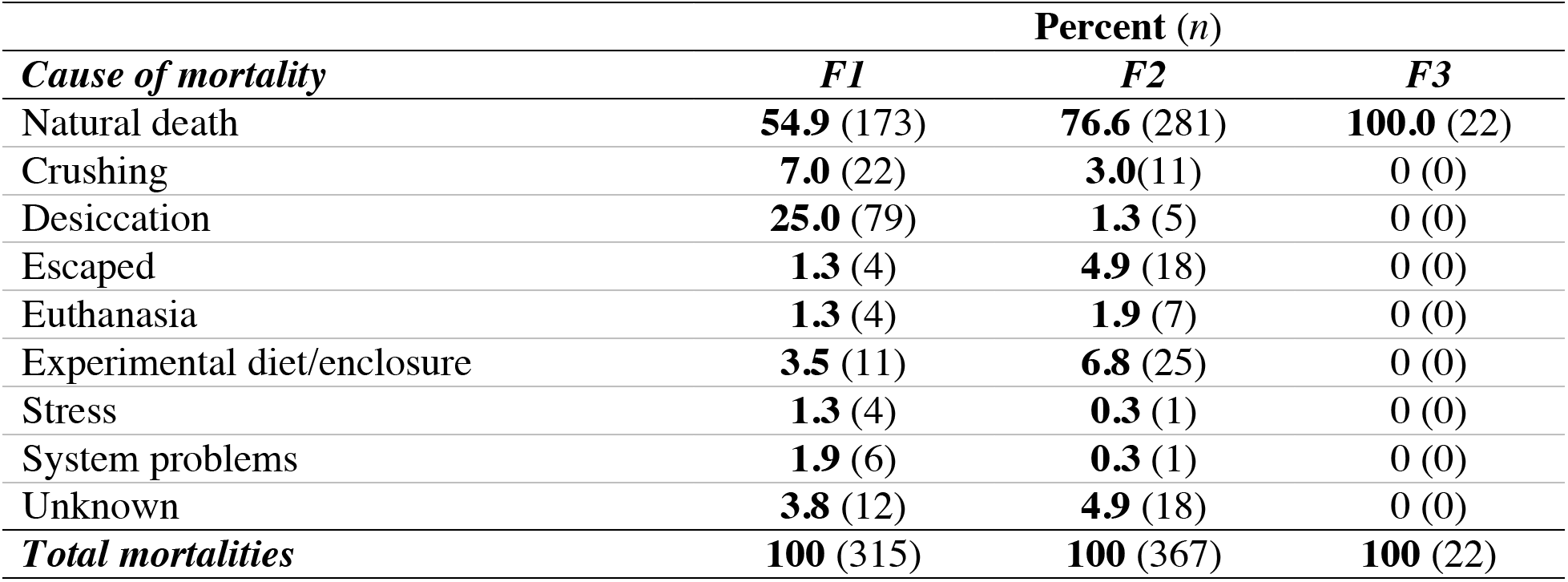
*O. chierchiae* mortalities by generation classified by cause of death. Natural deaths were defined as any mortality not caused by direct human interference or error and included death due to old age or any unknown health-related issues. Other non-natural causes of death included being crushed under enclosure lids (only occurred with the smallest hatchlings), escape from enclosures, euthanasia with 2-5% ethanol after observing signs of senescence or compromised health, use in experimental diets or enclosures, stress caused by external factors (such as poor water quality), system problems, or other unknown causes (i.e., cause of death was not recorded at time of mortality). F1=first generation; F2=second generation; F3=third generation*; n*=number of mortalities.

## 4. Discussion

Octopuses offer a unique and exceptional opportunity to compare complex nervous systems, behaviors, and development across evolutionary distant and anatomically divergent invertebrate and vertebrate model organisms. Studying octopuses from wild-collected specimens, however, restricts the scope of our understanding. New aquaculture methods are needed to rear octopuses in the laboratory. Here, we leveraged the small size and unique life cycle of *O. chierchiae* to establish the first multi-generational laboratory culture of the species. Starting with seven wild-collected individuals, we have been able to successfully rear *O. chierchiae* up to the third generation in large enough numbers to maintain a population spanning all life stages for over two years. All the data presented in this paper represents *O. chierchiae* behavior and life history of laboratory-bred populations. This study represents a significant advance for establishing these animals as a model system for biomedical research.

### 4.1 Size and growth considerations

Given their small body size, *O. chierchiae* require significantly less physical space to house compared to other octopus species. The maximum recorded mantle length of *O. chierchiae* in culture to date (40 mm) is dramatically smaller than other popular octopus model organisms, such as *O. bimaculoides, O. maya,* and *O. vulgaris*, which can reach sizes of up to 85 mm, 120 mm, and 250 mm in mantle length, respectively (Jereb et al., 2014). These species require much larger aquaria and occupy more space on a seawater system and physical space in a building (Wood et al., 2004), while *O. chierchiae* can be held in much smaller tanks. Previous studies have housed individual *O. bimaculoides* in 76 L tanks (Sinn et al., 2001) and *O. vulgaris* in 450 L tanks (Garcia and Valverde, 2006) while *O. chierchiae* can be housed in 2.84 L tanks. Based on reported adult enclosure sizes, 20 adults of the *O. chierchiae* can be held in a total of 56.8 L of seawater where *O. bimaculoides* would require 1520 L (Sinn et al., 2001) and *O. vulgaris* would require 9000 L of total seawater (Garcia and Valverde, 2006). Larger species of octopus also require more food and consequently produce more nitrogenous wastes (Boucher-Rodoni and Mangold, 1995; Oestmann et al., 1997; Lee et al., 1998). Culturing smaller octopuses like *O. chierchiae* thus reduces the biological load per individual on the seawater system, allowing them to be reared in closed seawater systems (Section 2.2), which are beneficial in maintaining consistent water chemistry and conditions when natural water chemistry in open seawater systems fluctuates.

*O. chierchiae* hatchlings and juveniles appear to display linear growth during the first 150-200 days of life (Figure 5). Based on the data from 40 randomly sampled adults between 280 to 800 days of age, average mantle length did not appear to increase with age after 280 days post-hatch (Figure 5). These results suggest that growth rates become asymptotic in adulthood, although this may be because diet was not increased during adulthood. Similarly, increasing temperature does not appear to increase growth rates past 6.5 months of age in *O. bimaculoides* (Forsythe and Hanlon, 1988). Average mantle length of adult *O. chierchiae* ranged from approximately 18-25 mm (Figure 5), and the largest mantle length observed was 40 mm. Growth rate and lifespan are known to be influenced by genetics and environmental factors in cephalopods. Higher temperatures influence egg development, accelerate growth rate early in life, reduce age at reproductive maturity, and shorten lifespan (Mangold and Boletzky, 1973; Forsythe and Hanlon, 1988; Forsythe et al., 1994; Forsythe et al., 2002 Vidal et al., 2002; Grigoriou and Richardson, 2008). Diet is known to increase the growth rate and size of several cultured octopus species (García and Valverde, 2006; Domingues et al., 2007; Rosas et al., 2007; Pham and Isidro, 2009; Prato et al., 2010; Baeza-Rojano et al., 2013). Future studies should investigate the effects of temperature and diet on *O. chierchiae* growth rates, reproduction, and lifespan.

### 4.2 Survival and lifespan

The mortalities that occurred in culture in this study have been largely due to natural causes while a minimal amount was due to human-related interactions (e.g., crushed under lids, experimental diets, direct handling) (Table 2). Reasons for natural death may include old age or other unknown health issues. Due to their low tolerance to ammonia (NH3) and nitrite (NO2), sources of nitrogenous waste in seawater systems should be removed as soon as possible (Oestmann et al., 1997; Lee et al., 1998). Rigorous water-quality management, frequent water changes, and regular maintenance of enclosures and seawater systems is critical for *O. chierchiae* culture. Frequent interactions from feeding, cleaning enclosures, and morphometric measurements may also be a source of stress if the octopuses are not handled gently. Natural deaths can be reduced through frequent water-quality checks combined with regular feeding and cleaning schedules, while minimizing the frequency of abrasive interactions with the octopuses. Over time, culture techniques of *O. chierchiae* at the Marine Biological Laboratory have greatly improved with continuous iterations of enclosure designs, feeding regimens, and maintenance techniques, along with rigorous training and attention to detail when working with hatchlings. This likely accounts for the decrease in “human-related” causes of death, such as human error, desiccation, and equipment malfunctions with each subsequent generation (Table 2).

In our study, mortality was highest in the first 30 days post-hatch across all generations, but cultured hatchlings regularly survived past 100 days (Figure 6), after which mortality decreased as they aged. Our data suggest that once *O. chierchiae* grow past a highly sensitive hatchling stage, they will be able to survive to sexual maturity. Our results are consistent with the low survival rate of paralarval octopus hatchlings in the wild (Laptikhovsky, 1998; Rocha et al., 2001) and a previous report of culturing *O. chierchiae* in which 4/26 hatchlings reached a maximum age of 67 days before dying (Rodaniche, 1984). This differs from published accounts of culturing other octopus species, in which mortality rates are steady over the entire life cycle (Hanlon and Forsythe, 1985). In our care, 14-15% of individuals in first- and second-generation clutches survived to over 400 days of age (Figure 6). Survival is lower in the third generation compared to first and second generations, although our present study is greatly limited by the COVID-19 pandemic (Figure 6). Typical signs of senescence exhibited by *O. chierchiae* at the end of life include reduced appetite, discoloration, and cessation of breeding (Anderson et al., 2002), but unlike other species of octopus, senescence is decoupled from the first incidence of reproduction (Rodaniche, 1984; Hofmeister et al., 2011). With further optimization of temperature or diet, cultured *O. chierchiae* may regularly live for up to two years and remain fertile for the majority of adulthood (Rodaniche, 1984). These results demonstrate that *O. chierchiae* has a comparable, if not longer, lifespan than many other octopus species commonly reared in culture, such as *O. bimaculoides* (1.5 years (Jereb et al., 2014)) and *O. maya* (300 days (Van Heukelm, 1977)). This would allow for observations and data collection on adult *O. chierchiae* for extensive periods of time while continuing to breed individuals to sustain the ongoing culture.

### 4.3 The advantage of iteroparity

Unlike most cephalopod species (Mangold, 1987; Cortez et al., 1995), *O. chierchiae* are iteroparous (Rodaniche, 1984; Hofmeister et al., 2011). Results from this study demonstrate that female *O. chierchiae* are capable of laying a clutch of eggs with mean size of 11-35 eggs approximately every 30 to 90 days while males are capable of mating several times without any apparent negative impact to their health. Some females laid multiple clutches of eggs after a single mating event, suggesting that they likely store sperm acquired from the male for an extended period of time. Sperm storage and sperm competition (Wigby and Chapman, 2004) has been previously reported in *O. chierchiae* (Rodaniche, 1984) and in other cephalopod species (Hanlon et al., 1999; Naud et al., 2005; Wada et al., 2005; Wada et al., 2010, Squires et al., 2015; Naud et al., 2016), although it is possible that sperm quality is reduced over time (Reinhardt, 2007). The rare iteroparous nature of *O. chierchiae* is extremely beneficial for laboratory culture; it affords multiple opportunities to harvest eggs at a distinct developmental stages and stagger age/size classes in culture.

Our data show that using the tasseling behavior (in males) or first incidence of egg-laying (in females) to estimate sexual maturity appeared to be sufficient for the breeding program to produce large numbers of hatchlings. In our study, the mean age at which females laid their first clutch of eggs was 241 days (Table 2). However, anecdotal necropsy observations performed during this study and from Rodaniche (1984) revealed the presence of eggs in females as young as 67 days of age. It is possible females may become sexually mature much earlier but do not lay eggs in the absence of sexually mature males. Based on the results from this study, we estimate that *O. chierchiae* reach sexual maturity at approximately six months of age, although timing of sexual maturation is likely modulated by temperature and diet. Future investigations are needed to determine the earliest laying time for this species and may be achieved by conducting mating trials at a younger age.

### 4.4 Applications in research

As J.Z. Young outlined in his classic work *<u>A Model of the Brain</u>*, octopuses offer a unique opportunity to discover ‘the rules’ of how to build complex brain function by comparing solutions across invertebrate and vertebrate nervous systems. Towards this goal, the advances in octopus husbandry we report here make *O. chierchiae* an especially attractive model organism in diverse scientific fields. Their predictable and repetitive reproduction provides a continuous supply of large, transparent eggs that permit access to the brain and other tissues using modern imaging techniques, such as calcium imaging, computed micro-tomography (micro-CT; Kerbl et al., 2013), ultrasound (Margheri et al., 2011), and magnetic resonance imaging (MRI; Tramacere et al., 2012; Jiang et al., 2014). This makes them an excellent candidate for the study of embryological development across different developmental stages, genetic lineages, and experimental groups. The large number of embryos produced by *O. chierchiae* make them an excellent octopus candidate for pioneering CRISPR-Cas9 gene-editing technology, which has been recently demonstrated in squid embryos (Crawford et al., 2020).

A steady supply of direct developing hatchlings provides many advantages to developmental, neurobiological, and behavioral study. Hatchlings exhibit many features of adults, such as hunting, which involves exquisite control of the arms and suckers. Since fluid forces vary across an animal’s size, strategies for controlling the flexible arms may change as the animal grows. A colony of *O. chierchiae* provides an opportunity to closely investigate the development and control of the chemotactile system of the arms across the entire lifespan, which has many applications in engineering and robotics. Octopuses also possess a camera-like eye and exhibit a range of visual behaviors, including hunting, predator evasion, navigation, and camouflage (Hanlon and Messenger, 2018). However, their brain organization is dramatically different from other highly visual animals such as vertebrates and insects, and little is known about the neural circuits that process visual information within the octopus brain. Standardized model species, particularly with genetic access, have played a key role in mechanistic studies of visual processing in other species, such as flies, zebrafish, and mouse (Huberman and Niell, 2011; Wernet, Huberman, Desplan 2014; Sanes and Zipursky 2010; Bollmann 2019), and the characteristics of *O. chierchiae* may allow it to serve this role for the unique visual system of cephalopods.

Iteroparity itself is interesting to study because it sets *O. chierchiae* apart from most other octopus species. In semelparous octopuses, reproduction triggers the signaling of the optic glands, the octopus analog of the vertebrate pituitary gland (Wang and Ragsdale 2018, Wells and Wells, 1969). The optic gland signaling systems are responsible for post-mating behavioral changes, including maternal behaviors and death, and have recently been characterized in *O. bimaculoides* for the first time (Wang and Ragsdale 2018). How the optic glands control reproduction in an iteroparous species is entirely unexplored. A lab-cultured colony of *O. chierchiae* would lay the foundation for investigating the interplay between reproduction and control of lifespan between iteroparous and semelparous octopus species, and accelerate the study of octopus reproductive neuroendocrinology, senescence, and aging.

Furthermore, *O. chierchiae* is closely related to the Larger Pacific Striped Octopus (LPSO), the only known social species of octopus (Caldwell et al., 2015). The LPSO is known to display unique behaviors such as beak-to-beak mating and mating pairs sharing a den (Caldwell et al., 2015). Importantly, some of the earliest insights into the brain mechanisms underlying social behaviors came from another such pair of asocial/social vertebrate sister species (montane and prairie voles) (Donaldson and Young, 2008). Unlike *O. chierchiae,* however, the LPSO is a small egg species and hatchlings undergo indirect development (Jereb et al., 2014; Caldwell et al., 2015). The existence of another comparator pair in invertebrates presents an enticing opportunity to test evolutionary hypotheses with the ultimate goal of discovering universal synaptic and circuit motifs that govern sociality across species.

### 4.5 *Current challenges to* O. chierchiae *culture*

There are still several challenges facing *O. chierchiae* culture. In addition to the careful attention to system design, feeding, and enclosure maintenance required to reduce mortality during the sensitive hatchling phase, our results suggest that the third generation began to experience the first signs of inbreeding depression (Charlesworth and Charlesworth, 1987). Although data from only one third-generation clutch was available, signs that point towards inbreeding depression in the third generation include: 1) significantly reduced survivorship, particularly in the first week of life (Figure 6), despite improved culture techniques and decreased mortality due to culture-related causes (Table 3); and 2) the birth of a 16-armed octopus, likely due to genetic mutations from the small gene pool. Given that our study began with only seven wild-collected individuals, this observation is not surprising. These findings emphasize the need to periodically supplement the population with new genetics via wild-collected octopuses. Decreased fertility in later generations of laboratory culture has been demonstrated in several cephalopod species including the common cuttlefish, *S. officinalis* (Forsythe et al., 1994), the flamboyant cuttlefish, *Metasepia pfefferi* (Grasse, 2014), and the bigfin reef squid, *Sepioteuthis lessoniana* (Walsh et al., 2002). More research on inbreeding depression and fecundity in *O. chierchiae* is required to help identify genetic bottlenecks and modify protocols to optimize breeding efficiency. Reduced survivorship observed in the third-generation clutch may also have been due to seasonal changes of water parameters of the incoming water source in open seawater systems, seasonal hormonal changes correlated to a breeding season, chemicals released by egg-laying females affecting the development of other eggs on the same system, or other unknown factors. Additional studies should explore these possibilities in order to understand *O. chierchiae* behavior, breeding, and biology. Due to the elusiveness of *O. chierchiae* in nature, researchers have been unable to study them in the ocean and little is known about their natural history in the wild. Future studies that investigate their distribution patterns, preferred habitats, natural diets, and typical reproductive behaviors in the wild will greatly improve laboratory husbandry practices.

### 4.6 Conclusion

*O. chierchiae* is an ideal model organism due to its small size, iteroparity, and predictable reproduction, and direct developing hatchlings. Various aspects of octopus behavior and biology make laboratory culture challenging: a rapid metabolism that emphasizes the need for optimal water quality and frequent feeds, an affinity to escape enclosures, cannibalistic tendencies, and the requirement for large amounts of physical space. We have developed a culture program for *O. chierchiae* that optimizes reproductive success and survivorship to address the various challenges of octopus aquaculture. Starting with seven wild-collected individuals, we successfully reared *O. chierchiae* up to the third generation in large enough numbers to maintain a population spanning all life stages for over two years. More research in both the laboratory and the wild is needed to investigate the effects of temperature and diet on growth rates, reproduction, and lifespan, to identify the precise timing and cues for the onset of sexual maturity in this species, and to study mating behaviors in order to optimize breeding programs and maximize reproductive output. In the long term, periodically supplementing an *O. chierchiae* culture with new genetic pools from wild-collected octopuses likely would reduce inbreeding-related defects and mortality, which will be important for maintaining a healthy, sustainable culture of *O. chierchiae.*

## Ethical note

In the USA, cephalopods are not included in federal regulations that govern the use of animals in research laboratories, thus no protocol or approval number was required for this study. However, the Marine Biological Laboratory voluntarily upholds a Cephalopod Care Policy and the care of the animals in this study adhered to that policy. All animal care and handling followed methods established by the regulations of the Directive 2010/63/EU for cephalopods (Fiorito et al., 2015a; Smith et al., 2013; Sykes et al., 2012) and institutional standards for cephalopod care.

## Author contribution statement

AGG, AD, TS, and BG contributed to the conception and design of the study. AGG, AD, and ZYW analyzed the data and wrote the manuscript. All authors contributed to manuscript revision, read, and approved the submitted version.

## Acknowledgements

We acknowledge the pioneering culture efforts of Rich Ross, Danielle Dallis, Travis Snyder, Benjamin Liu, Anna Ramji, Leo Song, and Saumitra Kelkar; Mathew Everett, Peter Kilian, and all former interns in the Cephalopod Program at the Marine Biological Laboratory (MBL) for assistance with data collection and ongoing *O. chierchiae* culture; and Philip Swiny, Carlos Carstens, Juan-chi, Vanessa Grasse, and William Wcislo (Smithsonian Tropical Research Institute) for support with field collection investigations. We thank Roger Hanlon, Jonathon Stone, and David Welch for providing feedback on earlier versions of the manuscript, Tim Briggs for his exceptional photographs of the octopuses, and Caroline Albertin for research support. We are grateful for Joshua Rosenthal’s support of the MBL’s cephalopod culture efforts, and for John Sigel and Sally Reid’s support of the Northeastern University Co-op Internship Program at the MBL.

