## Supplementary Figures and Legends for "The lesser Pacific striped octopus, *Octopus chierchiae*: an emerging laboratory model for the study of octopuses"

**Supplementary material**

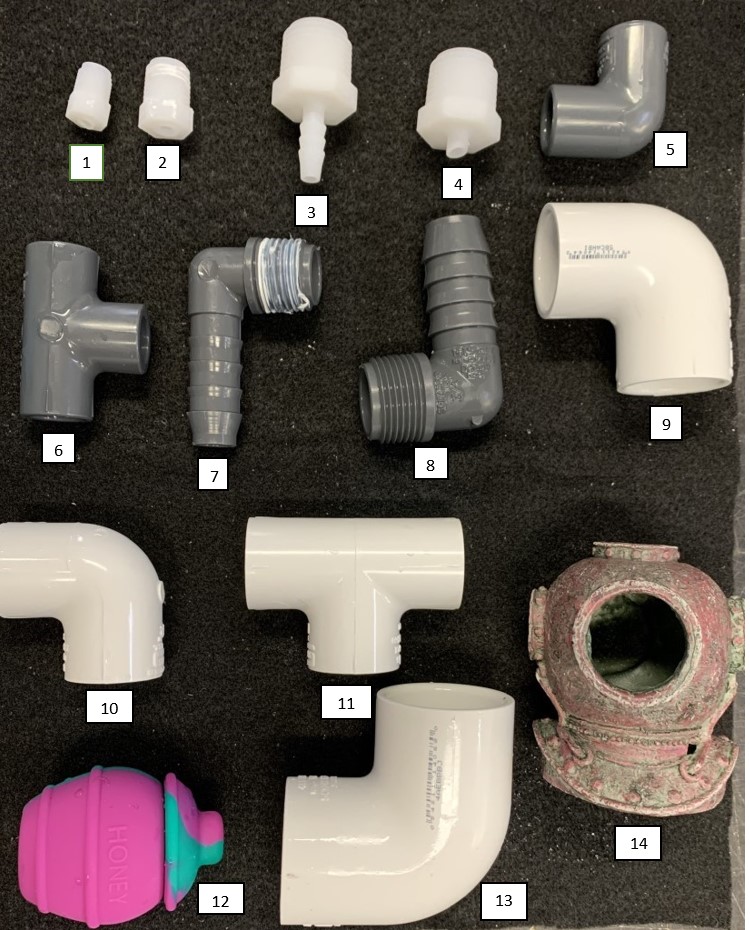

**Figure S1.** **Den options.** Dens provided in hatchling enclosures (1-2), juvenile enclosures (3-7), and adult enclosures (7-14).

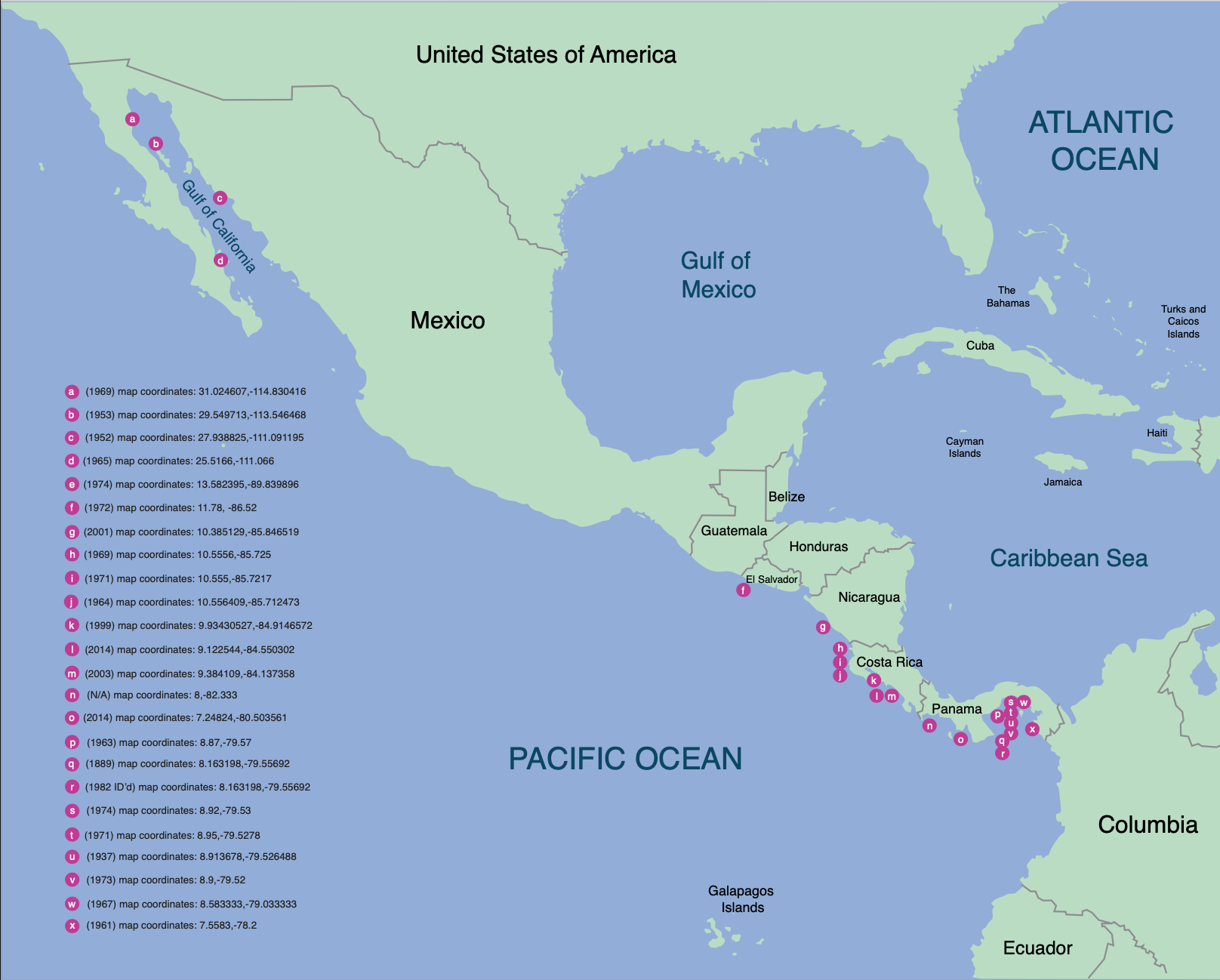

**Figure S2. Map of *O. chierchiae sightings* from 1889 to 2014.** Locations where *O. chierchiae* have been sighted/collected from 1889 to 2014. References for each location are available in Table S2.

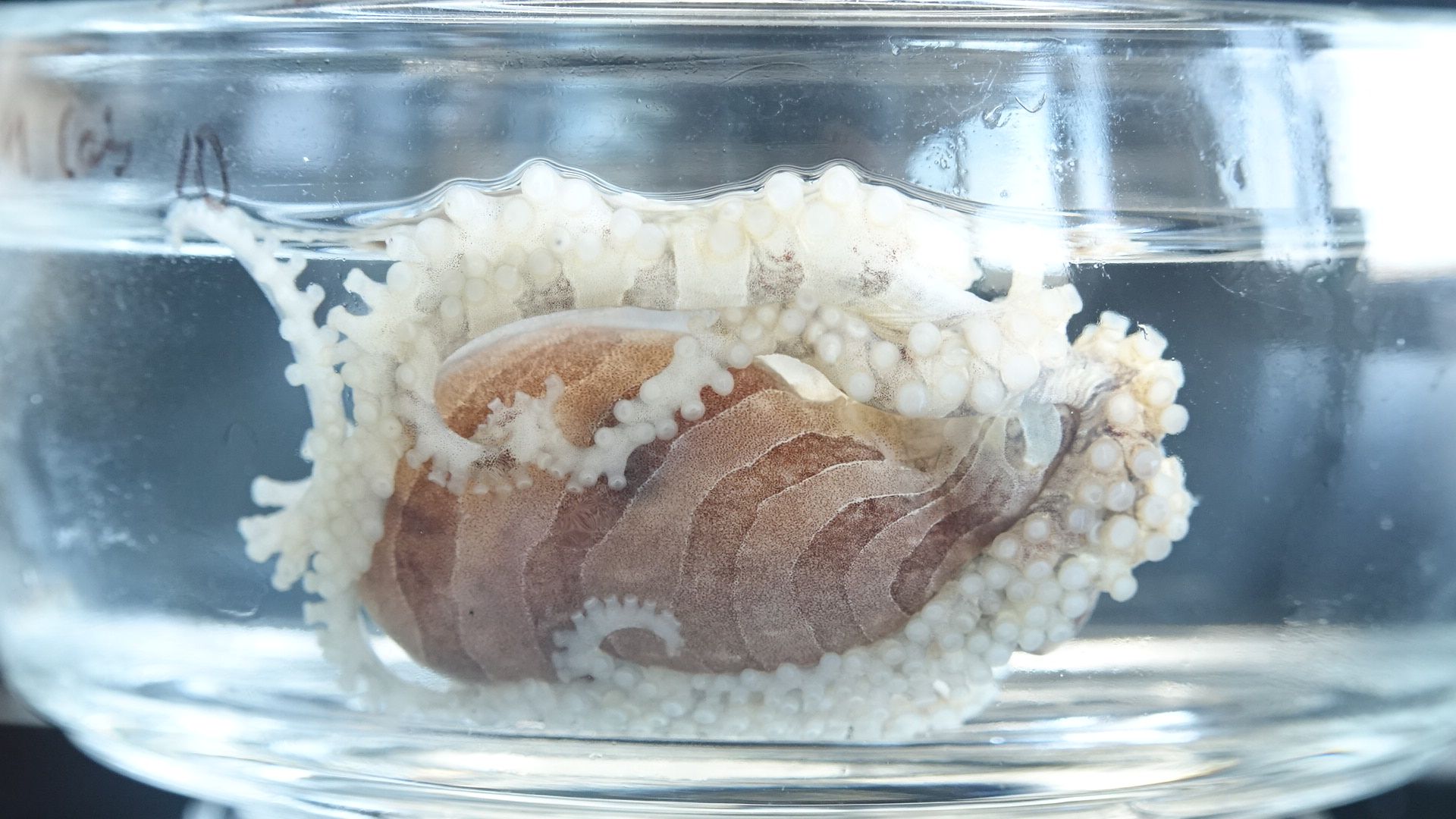

**Figure S3. Adult female *O. chierchiae* exhibiting signs of senescenc*e*.** Typical signs of octopus senescence (Anderson et al., 2002) include disorientation, inappetence, discoloration, and abnormal locomotion, all of which are found in senescing *O. chierchiae.* This female exhibits an inverted resting posture, which is atypical of healthy adults, and increased lethargy. Individuals ceased reproduction after onset of senescence.

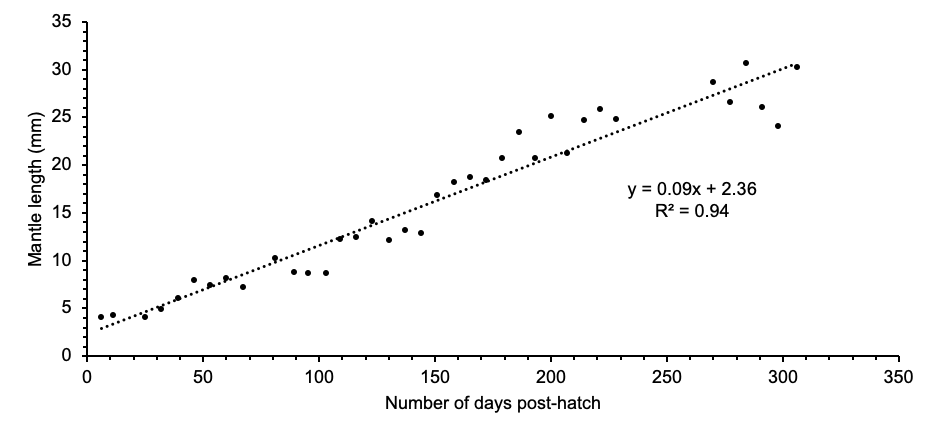

**Figure S4. Mantle length of an individual *O. chierchiae* over time.** To minimize handling stress, photographs were taken within a minute of moving the octopus to the the measuring chamber, and fluctuation in mantle length is partly due to subtle differences in the octopus’ posture at time of photography. Each point represents a separate measurement (*n*=37).

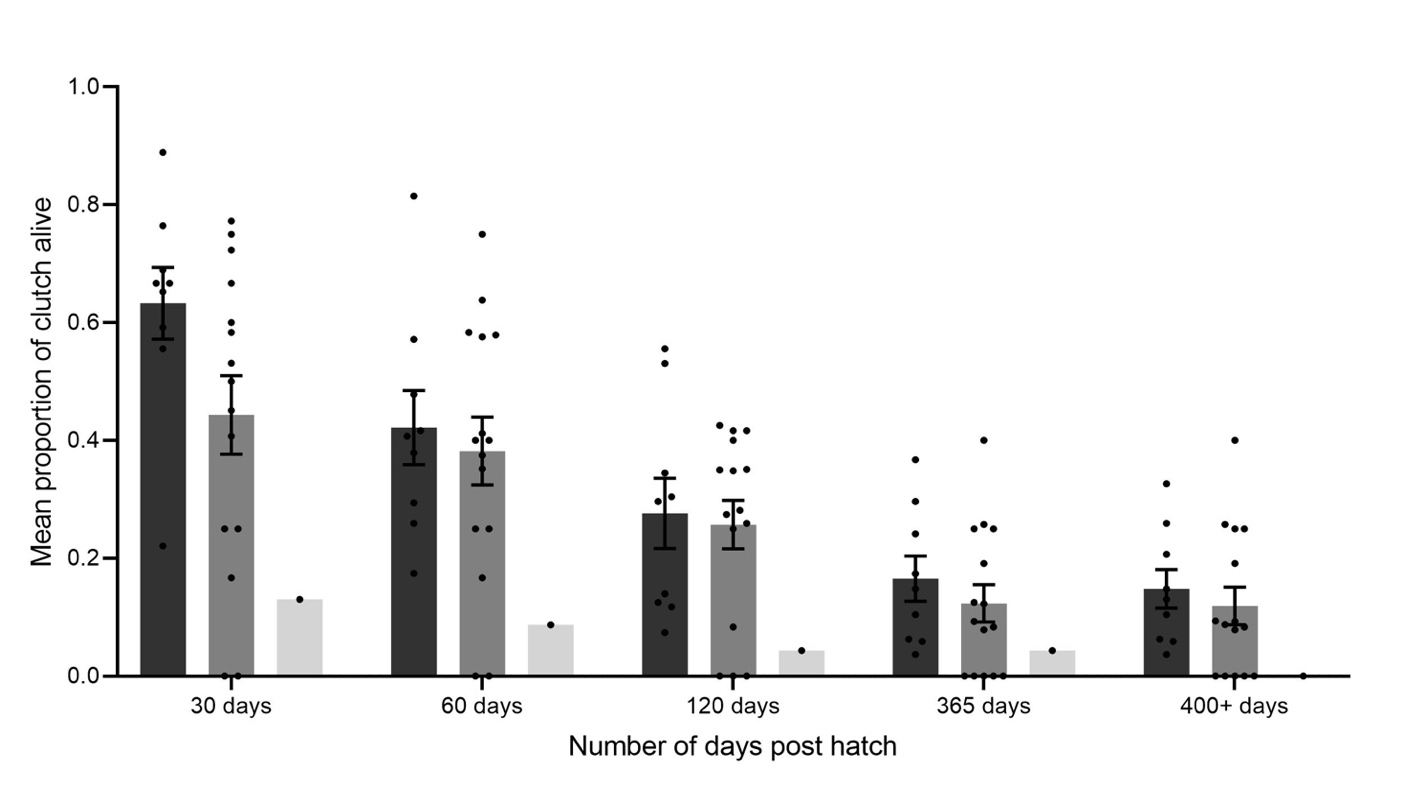

**Figure S5. Proportion of *O. chierchiae* clutch surviving over** **time.** Mean proportion (±SEM) of survivors per clutch over time. A two-way analysis of variance followed by Tukey’s multiple comparison test revealed that the proportion of survivors between first and second generation was not significantly different across all time points. The total proportion of survivors for the single third-generation clutch is reported. F1=first generation; F2=second generation; F3=third generation; *n*=number of clutches; SEM=standard error of the mean.

**See attached files for the following:**

**Table S1. Reproductive mating pairs.** Mating pairs that produced viable clutches of eggs, sorted by the dam (i.e., mother). A total of 15 females and seven males reproduced. Eight females laid more than one clutch of eggs and five males sired several clutches. One male (R(18)11), sired eight different clutches for a total of 214 hatchlings. Most of the wild females (WF) had mated previously with unknown partner(s) and produced one or more clutches upon arrival in the lab.

**Table S2. Location of *O. chierchiae* sightings from 1889 to 2014.**

**Supplementary Video 1. Male *O. chierchiae tasseling*** – Male *O. chierchiae* exhibiting tasseling behavior during a mating trial**.** This behavior was used as an indirect proxy to estimate sexual maturity in males.

**Supplementary Video 2. Mating trial (full duration)** *–* Full mating trial.

**Supplementary Video 3. Mating trial (close-up)** – First 45 seconds of a mating trial, showing the male pouncing on the female and the male’s hectocotylized arm searching for female’s mantle opening.
